# The high-affinity immunoglobulin receptor FcγRI potentiates HIV-1 neutralization via antibodies against the gp41 N-heptad repeat

**DOI:** 10.1101/2020.08.27.271064

**Authors:** David C. Montefiori, Maria V. Filsinger Interrante, Benjamin N. Bell, Adonis A. Rubio, Joseph G. Joyce, John W. Shiver, Celia C. LaBranche, Peter S. Kim

**Author notes:** co-first authors (David C. Montefiori, Maria V. Filsinger Interrante).

## Abstract

The HIV-1 gp41 N-heptad repeat (NHR) region of the pre-hairpin intermediate, which is transiently exposed during HIV-1 viral membrane fusion, is a validated clinical target in humans and is inhibited by the FDA-approved drug enfuvirtide. However, vaccine candidates targeting the NHR have yielded only modest neutralization activities in animals; this inhibition has been largely restricted to tier-1 viruses, which are most sensitive to neutralization by sera from HIV-1-infected individuals. Here, we show that the neutralization activity of the well-characterized NHR-targeting antibody D5 is potentiated >5,000-fold in TZM-bl cells expressing FcγRI compared to those without, resulting in neutralization of many tier-2 viruses (which are less susceptible to neutralization by sera from HIV-1-infected individuals and are the target of current antibody-based vaccine efforts). Further, antisera from guinea pigs immunized with the NHR-based vaccine candidate (ccIZN36)_3_ neutralized tier-2 viruses from multiple clades in an FcγRI-dependent manner. As FcγRI is expressed on macrophages and dendritic cells, which are present at mucosal surfaces and are implicated in the early establishment of HIV-1 infection following sexual transmission, these results may be important in the development of a prophylactic HIV-1 vaccine.

## Introduction

Membrane fusion between HIV-1 and host cells is mediated by the viral envelope glycoprotein (Env), a trimer consisting of the gp120 and gp41 subunits. Upon interaction with cellular receptors, Env undergoes a dramatic conformational change and forms the pre-hairpin intermediate (PHI) (1–3), in which the fusion peptide at the amino terminus of gp41 inserts into the cell membrane. In the PHI, the N-heptad repeat (NHR) region of gp41 is exposed and forms a stable, three-stranded a-helical coiled coil. Subsequently, the PHI resolves when the NHR and the C-heptad repeat (CHR) regions of gp41 associate to form a trimer-of-hairpins structure that brings the viral and cell membranes into proximity, facilitating membrane fusion (Figure 1).

**Figure 1.**
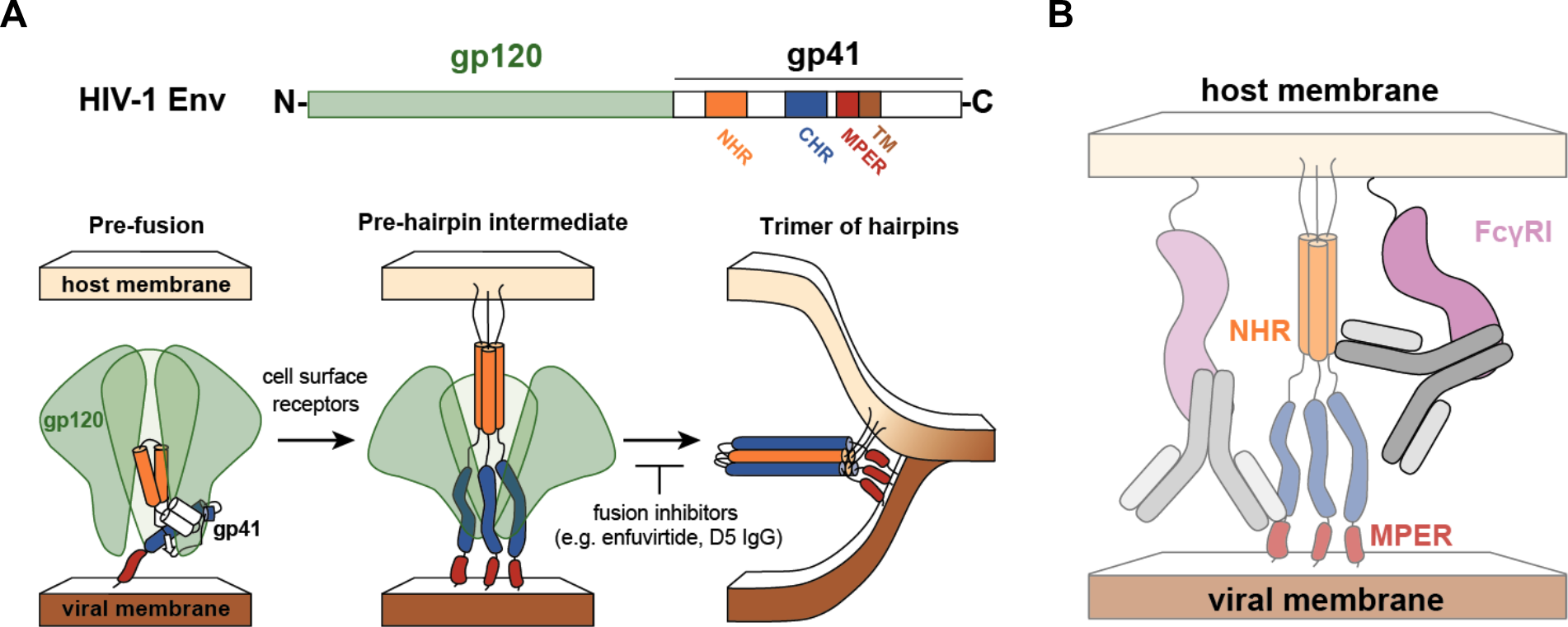
HIV-1 membrane fusion and potentiation by FcγRI. **(A)** The surface protein of HIV-1, Envelope (Env), is composed of the gp120 and gp41 subunits. After Env binds to cell-surface receptors, gp41 inserts into the host cell membrane and undergoes a conformational change to form the pre-hairpin intermediate (PHI). The N-heptad repeat (NHR) region of gp41 is exposed in the PHI and forms a three-stranded coiled coil. To complete viral fusion, the PHI resolves to a trimer-of-hairpins structure in which the C-heptad repeat (CHR) adopts a helical conformation and binds the NHR region. Fusion inhibitors such as enfuvirtide bind the NHR, preventing viral fusion by inhibiting formation of the trimer-of-hairpins (1–3). The MPER is located adjacent to the transmembrane (TM) region of gp41. **(B)** Hypothesized mechanism for FcγRI-mediated potentiation of antibodies targeting the NHR. The Fc domain of the antibody is bound by FcγRI, similar to the previously characterized mechanism of FcγRI-mediated potentiation of antibodies targeting the MPER (17, 18). Both the NHR and the MPER are inaccessible or only partially accessible in the native state of Env.

The NHR region of the PHI is a validated therapeutic target in humans: the FDA-approved drug enfuvirtide binds the NHR and inhibits viral entry into cells (4, 5). Various versions of the three-stranded coiled coil formed by the NHR have been created and used as vaccine candidates in animals (6–10). The neutralization potencies of these antisera, as well as those of anti-NHR monoclonal antibodies (mAbs) (11–15), are modest and mostly limited to HIV-1 isolates that are highly sensitive to antibody-mediated neutralization (commonly referred to as tier-1 viruses (16)). These results have led to skepticism about the PHI as a vaccine target.

Earlier studies showed that the neutralization activities of mAbs that bound another region of gp41, the membrane proximal external region (MPER), were enhanced as much as 5,000-fold in cells expressing FcγRI (CD64), an integral membrane protein that interacts with the Fc portion of IgG molecules (17). This effect was not attributed to phagocytosis and occurred when the cells were preincubated with antibody and washed before adding virus (17, 18). Since the MPER is a partially cryptic epitope that is not fully exposed until after Env engages with cellular receptors (19, 20), these results suggest that by binding the Fc region of MPER mAbs, FcγRI provides a local concentration advantage at the cell surface that enhances viral neutralization (17, 18). While not expressed on T cells, FcγRI is expressed on macrophages and dendritic cells (21), which are present at mucosal surfaces and are implicated in sexual HIV-1 transmission and the early establishment of HIV-1 infection (22–34).

Here we investigated whether FcγRI expression also potentiates the neutralizing activity of antibodies targeting the NHR, since that region, like the MPER, is preferentially exposed during viral fusion. We found that D5, a well-characterized anti-NHR mAb (11, 12), inhibits HIV-1 infection ~5,000-fold more potently in TZM-bl cells expressing FcγRI (TZM-bl/FcγRI cells) than in TZM-bl cells that do not. Further, while antisera from guinea pigs immunized with (ccIZN36)_3_, an NHR-based vaccine candidate (7), displayed weak neutralizing activity in TZM-bl cells, they exhibited enhanced neutralization in TZM-bl/FcγRI cells, including against some tier-2 HIV-1 isolates that are more resistant to antibody-mediated neutralization (16) and that serve as benchmarks for antibody-based vaccine efforts. These results indicate that FcγRI can play an important role in neutralization by antibodies that target the PHI. Since these receptors are expressed on cells prevalent at mucosal surfaces thought to be important for sexual HIV-1 transmission, our results motivate vaccine strategies that harness this potentiating effect.

## Results

D5, a mAb shown by X-ray crystallography to bind a highly conserved epitope on the NHR (12), has weak but relatively broad neutralizing activity against HIV-1 strains (11). We measured the neutralizing activity of D5 against HXB2 (a clade B tier-1 virus) in both TZM-bl and TZM-bl/FcγRI cells. Notably, the presence of FcγRI increased the neutralization potency of D5 by approximately 5,000-fold (Figure 2A). In contrast, enhanced neutralization in TZM-bl/FcγRI cells was not observed with the Fab fragment of D5 (Figure 2A), indicating that this phenomenon is Fc-dependent. In addition, this enhancement was not detected when D5 IgG was tested in TZM-bl cell lines expressing FcγRIIa, FcγRIIb, or FcγRIIIa (Figure 2B), suggesting that the observed effect is specific to FcγRI.

**Figure 2:**
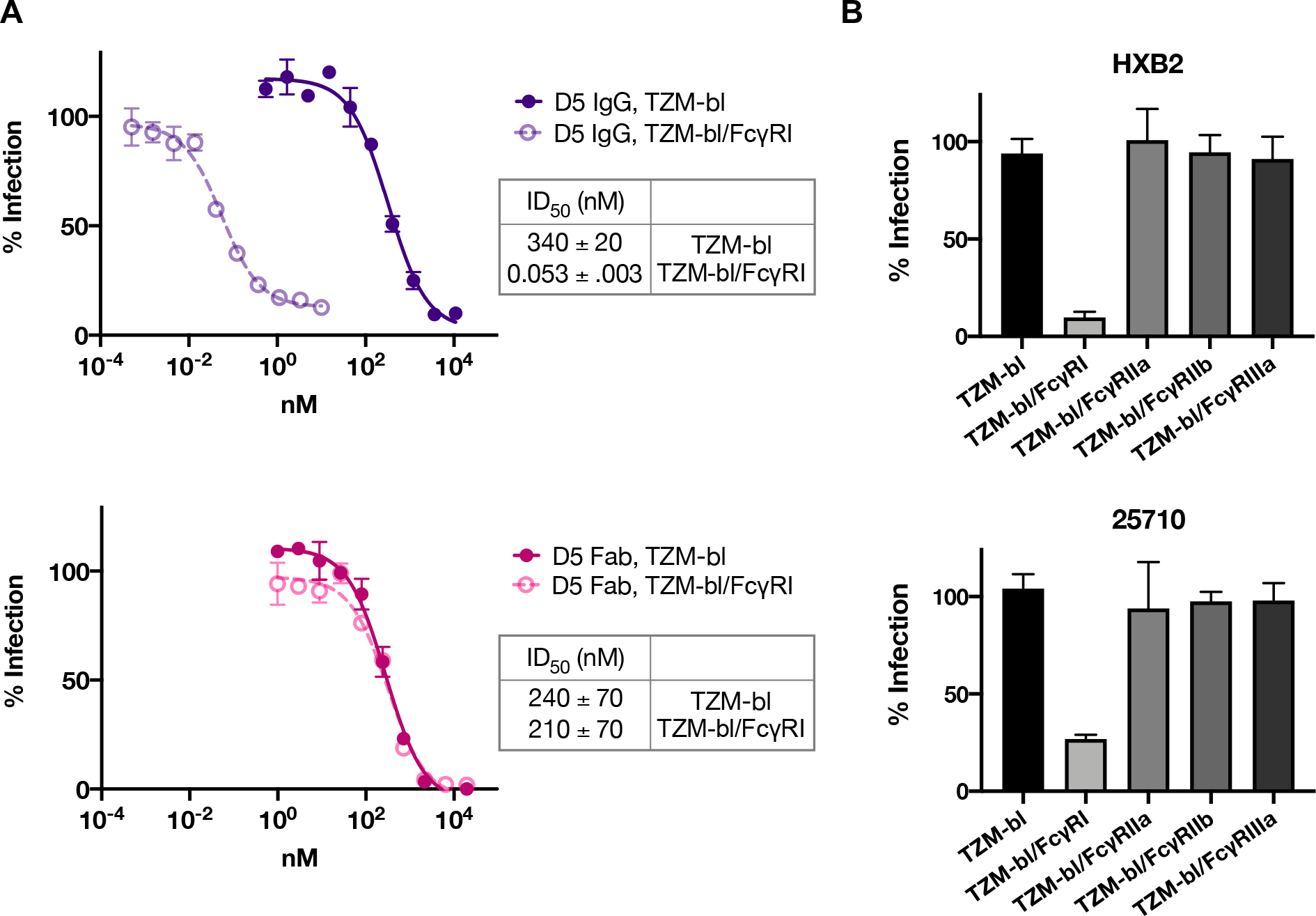
The neutralization potency of the anti-NHR antibody D5 is enhanced by FcγRI. **(A)** Inhibition of infection by viruses pseudotyped with Env from HXB2 (tier 1, clade B) by D5 IgG (top) and D5 Fab (bottom) in TZM-bl cells not expressing (solid) or expressing (open) FcγRI. Potentiation of >5,000-fold occurs in TZM-bl/FcγRI cells for the IgG but not the Fab form of D5. Curves plotted are from a single experiment; error bars are the range of *n*=2 measurements. The table shows ID_50_ (50% inhibitory dose) mean values and standard error of the mean (SEM) from duplicate experiments. **(B)** D5 IgG (0.5 μg/mL) inhibition of infection of Env-pseudotyped lentiviruses (top, HXB2; bottom, 25710) in TZM-bl cells stably expressing various Fcγ receptors. Mean and standard deviation were plotted for *n*=8 replicates. The observed potentiation effect is specific to FcγRI in both viruses tested.

D5 weakly inhibits a diverse range of HIV-1 viruses across clades, as expected given the high (>95%) conservation of residues that form the D5 epitope on the NHR (11, 12). Given the increase in potency afforded by FcγRI, we investigated the neutralization by D5 in a wide panel of HIV-1 pseudotyped viruses (35) in TZM-bl/FcγRI cells. At concentrations up to 25 μg/mL, D5 failed to neutralize viruses in the panel when measured in TZM-bl cells not expressing FcγRI (Table 1). However, when measured across the same concentration range in TZM-bl/FcγRI cells, D5 inhibited eight of the nine tier-2 viruses in the panel, spanning five clades (Table 1).

**Table 1:** D5 IgG neutralizes tier-1 and tier-2 viruses across clades in TZM-bl/FcγRI cells. Viruses were pseudotyped with Env from various HIV-1 strains. 50% inhibitory dose (ID_50_) was determined using a validated neutralization assay (49–52). SVA-MLV is lentivirus pseudotyped with murine leukemia virus (MLV) envelope to detect non-specific inhibition.

| Virus | | | ID<sub>50</sub> in<br>TZM-bl cells<br>($\mu\text{g/mL}$) | ID<sub>50</sub> in<br>TZM-bl/FcγRI cells<br>($\mu\text{g/mL}$) |
| --- | --- | --- | --- | --- |
| **SVA-MLV** | neg. ctrl. |  | >25 | >25 |
| **X2278** | Tier 1B | Clade B | >25 | 0.53 |
| **246-F3** | Tier 2 | Clade AC | >25 | >25 |
| **CNE55** | Tier 2 | CRF01_AE | >25 | 0.88 |
| **TRO.11** | Tier 2 | Clade B | >25 | 4.8 |
| **BJOX2000** | Tier 2 | CRF07_BC | >25 | 0.53 |
| **CH119** | Tier 2 | CRF07_BC | >25 | 1.8 |
| **Ce1176** | Tier 2 | Clade C | >25 | 7.0 |
| **25710** | Tier 2 | Clade C | >25 | 0.36 |
| **Ce0217** | Tier 2 | Clade C | >25 | 0.43 |
| **X1632** | Tier 2 | Clade G | >25 | 0.71 |

Consistent with earlier studies (17), addition of normal human serum to the neutralization assay diminished the potentiation of D5 neutralizing activity in TZM-bl/FcγRI cells (Figure 3). This decrease in neutralization was dependent on the concentration of added human serum (Figure 3), as expected if serum IgG competes for binding to FcγRI and thereby reduces the potentiation of D5 neutralizing activity. Because D5 has such high potency in TZM-bl/FcγRI cells (ID_50_ < 0.01 μg/mL, Figure 2A), it is not surprising that 0.5% human serum (~50 μg/mL IgG) greatly diminishes the observed potentiation (Figure 3). However, in a vaccine setting, where a substantial fraction (~1% to ~10%) of serum immunoglobulin can be antigen specific (36–39), antisera containing vaccine-elicited antibodies against the NHR may exhibit FcγRI-dependent neutralization enhancement. Indeed, antisera from guinea pigs immunized with the NHR-based vaccine candidate (ccIZN36)_3_, which had weak or no neutralizing activity in TZM-bl cells, neutralized tier-2 viruses from multiple clades when tested in TZM-bl/FcγRI cells (Figure 4). These data demonstrate that even in the presence of non-vaccine-elicited serum IgG, vaccine-elicited antibodies against the NHR have enhanced FcγRI-dependent neutralization.

**Figure 3:**
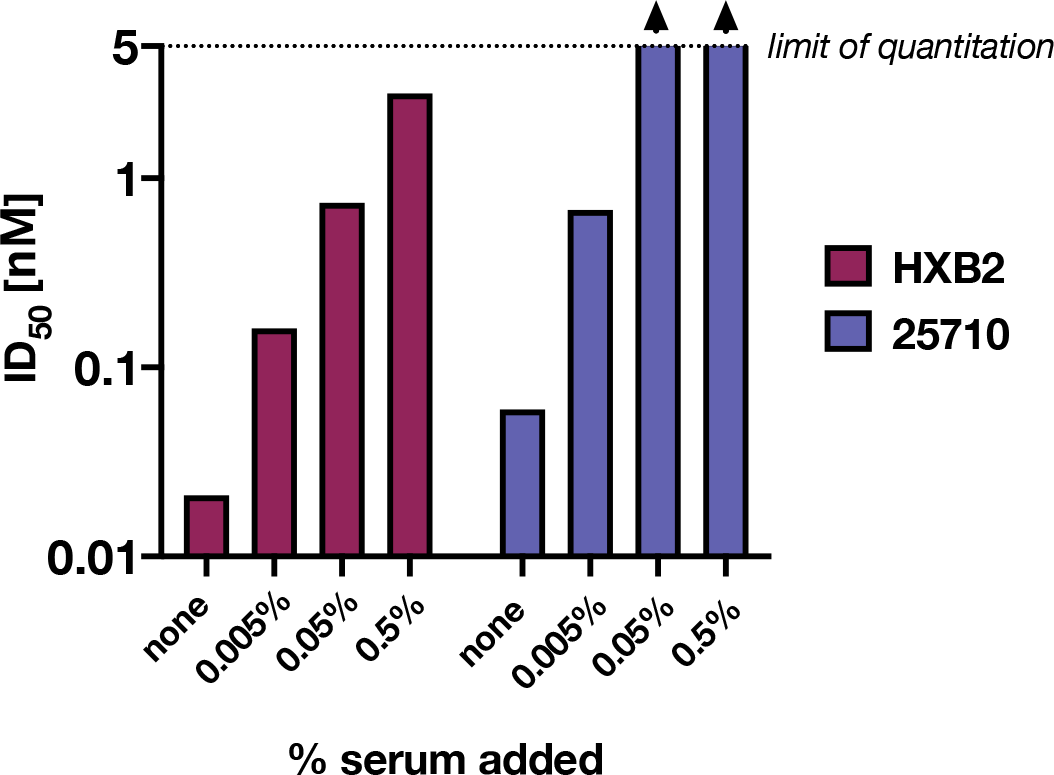
Addition of normal human serum to infection assays diminishes the potentiation of D5 neutralization activity by FcγRI. Values of 50% inhibitory dose (ID_50_) for D5 IgG (nM) against viruses pseudotyped with Env from two HIV-1 strains (HXB2 and 25710) measured in the presence of 0.005% to 0.5% added human serum. Values above the limit of quantitation for this assay (5 nM) are indicated with arrows. ID_50_ values were obtained from non-constrained fits of 4- and 5-point dilution curves, where the neutralization value at each dilution was measured in duplicate.

**Figure 4:**
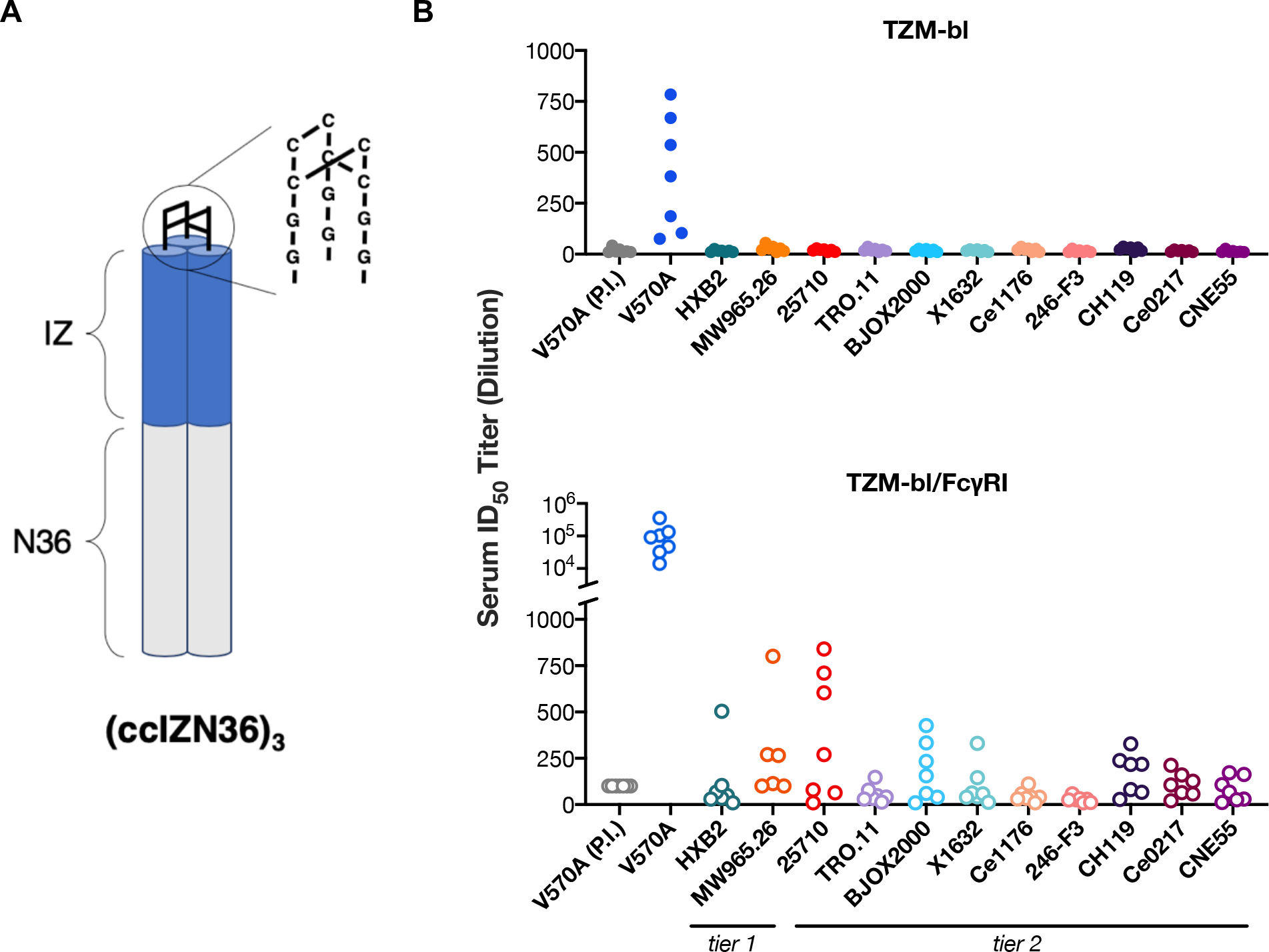
Antisera from guinea pigs immunized with an NHR-based vaccine candidate, (ccIZN36)_3_, neutralize multiple tier-2 HIV-1 strains in TZM-bl/FcγRI cells. **(A)** (ccIZN36)_3_ contains 36 residues from the NHR region of gp41 (HXB2) and is stabilized by disulfide bonds and a coiled-coil domain (7). **(B)** 50% inhibitory dose (ID_50_) titers (serum dilution) were determined for antisera against viruses pseudotyped with Env from various HIV-1 strains measured in TZM-bl cells not expressing (top, closed) or expressing (bottom, open) FcγRI using a validated neutralization assay (49–52). V570A is a mutant of the HXB2 strain that is more sensitive to antibodies that target the PHI (7), and (P.I.) denotes pre-immune antisera tested against this strain. Pre-immune antiserum from each animal was tested against all viruses and did not have detectable neutralization in any of the strains (ID_50_ titers below the limit of quantitation, which was 100 for TZM-bl/FcγRI assays with V570A antisera and 10 for all others; data not shown). Each data point represents the ID_50_ value of antiserum from a single guinea pig after a prime and two boosts with (ccIZN36)_3_.

## Discussion

Here we have established that D5, a mAb targeting the NHR region of the PHI, is ~5,000-fold more potent at preventing HIV-1 infection in FcγRI-expressing cells than in cells that do not express this receptor. Earlier studies reported a similar effect with MPER-binding mAbs, including 2F5, 4E10 (17, 18), and 10E8 (L.G. Perez & D.C. Montefiori, unpublished results), but not with mAbs that target other HIV-1 epitopes. Importantly, these earlier studies also revealed that this enhancement occurred when cells were preincubated with antibody and washed before virus was added.

Taken together, these results support a model for potentiation in which prepositioning of antibodies by FcγRI at cell surfaces increases the local concentration of antibodies and thereby enhances neutralization (17, 18). Such a mechanism would be expected to impact HIV-1 antibodies that target epitopes on Env that are only exposed after engagement with cellular receptors, such as the MPER or the NHR. Since other viruses that utilize type I fusion proteins appear to proceed through a PHI during cell entry (2, 3), potentiation of anti-PHI antibodies against other viruses can also be expected.

Although not normally expressed on T cells, FcγRI is expressed on macrophages and dendritic cells (21), which are present at the mucosal surfaces where HIV-1 transmission occurs. Both macrophages and dendritic cells can be productively infected by HIV-1 (22–25) and transmit virus to CD4+ T cells (26–29). Studies of intravaginal SIV inoculation of non-human primates demonstrate infection of intraepithelial and submucosal dendritic cells at the earliest stages of infection (30–32). Importantly, the migration of HIV-1 positive dendritic cells from the initial site of infection to lymph nodes results in dissemination of virus to large numbers of CD4+ T cells (32–34). Thus, it is plausible that inhibition of HIV-1 infection of FcγRI-expressing cells could decrease the likelihood of HIV-1 transmission.

Two previous studies of non-human primates support the notion that FcγRI may have an important role in protection provided by MPER antibodies against simian-human immunodeficiency virus (SHIV) challenge. First, in a vaginal challenge with SHIV-BaL in rhesus macaques, dose-dependent protection was observed for an MPER mAb (2F5) when it was administered as an IgG, but not with its Fab form, despite higher vaginal Fab levels at the time of challenge (40), implicating an Fc-dependent mechanism. Second, in a comprehensive meta-analysis of passive immunization studies showing that serum-neutralization antibody titers associate with protection against SHIV challenge, mAbs targeting the MPER were a highly significant outlier compared to other neutralizing mAbs, with much greater potency than would be predicted from serum-neutralization titers measured in cell culture (41). These findings motivate efforts to investigate the ability of passively transferred anti-PHI antibodies to protect against sexual HIV-1 transmission.

Finally, here we demonstrate that antisera elicited with a vaccine candidate against the NHR of the HIV-1 PHI show substantially enhanced neutralization activity mediated by FcγRI. In particular, antisera from guinea pigs immunized with (ccIZN36)_3_ exhibited cross-clade neutralization of a diverse panel of tier-2 viruses in an FcγRI-dependent manner (Figure 4). Interrogating the role of FcγRI potentiation of neutralization *in vivo* and the impact of anti-NHR antibodies in protection are therefore important lines of inquiry in the development of an effective HIV-1 vaccine.

## Materials and Methods

### Antibody expression and purification

D5 IgG and Fab were produced in Expi293F cells. Constructs were cloned using the In-Fusion® HD Cloning Kit master mix (Clontech); the heavy and light chain regions were cloned into the CMV/R plasmid backbone for expression under a CMV promoter. This vector includes the HVM06_Mouse (P01750) Ig heavy chain V region 102 signal peptide to induce protein secretion and to enable purification from the supernatant. These plasmids were transfected into Expi293F cells at 3×10^6^ cells/mL with FectoPRO® (Polyplus), with the antibody heavy and light chain plasmids co-transfected at a 1:1 ratio. Cell cultures were incubated at 37 °C and 8% CO2 with shaking at 120 rpm on a MaxQ^™^ 2000 CO2-resistant digital shaker (Thermo Fisher Scientific). Cells were harvested 3 days post transfection by spinning at 4000 x *g* for 15 min and filtered through a 0.22-μm filter. IgG-containing supernatants were diluted 1:1 with 1x phosphate-buffered saline [pH 7.4] and batch-bound to Pierce™ Protein A agarose (Thermo Fisher Scientific) overnight at 4 °C. The supernatant/resin slurry was added to a column and the resin was washed with 1x phosphate-buffered saline [pH 7.4] and eluted with 100 mM glycine [pH 2.8] into 1/10 volume of 1 M Tris [pH 8.0]. Similarly, Fab-containing supernatants were diluted 1:1 with 50 mM sodium acetate [pH 5.0], batch-bound to Pierce™ Protein G Agarose (Thermo Fisher Scientific) overnight at 4 °C, washed with 50 mM sodium acetate [pH 5.0], and eluted with 100 mM glycine [pH 2.8] into 1/10 volume of 1 M Tris [pH 8.0].

### Viral neutralization assay

Neutralizing antibody activity of monoclonal antibodies and serum samples was measured in 96-well culture plates using Tat-regulated luciferase reporter gene expression to quantify reductions in virus infection in TZM-bl and TZM-bl/FcγRI cells. TZM-bl cells were obtained from the NIH AIDS Research and Reference Reagent Program, as contributed by John Kappes and Xiaoyun Wu (42–46). TZM-bl cells transduced to stably express FcγRI, TZM-bl/FcγRI (17, 18), were also used as target cells in the neutralization assays.

Neutralization assays were performed using well-established Env-pseudotyped lentiviral reference strains (16, 35, 47, 48) in TZM-bl and TZM-bl/FcγRI cells essentially as previously described (49). Serum samples were heat-inactivated at 56 °C for 1 h, then diluted over a range of 1:20 to 1:43740 in cell culture medium and pre-incubated with virus (~150,000 relative light unit equivalents) for 1 h at 37 °C before addition of cells. Experiments with D5 IgG reported in Table 1 included the 1 h 37 °C incubation, while those in Figure 2 and 3 did not. After incubation for 48 h, cells were lysed and luciferase activity was determined using a microtiter plate luminometer and BriteLite Plus Reagent (Perkin Elmer). Neutralization titers were defined as the sample dilution at which relative luminescence units (RLU) were reduced by 50% compared to RLU in virus control wells after subtraction of background RLU in cell control wells. This assay was previously optimized and validated (49, 50) and was conducted in compliance with good clinical laboratory procedures (51), including participation in a formal TZM-bl assay proficiency program for GCLP-compliant laboratories (52).

### (ccIZN36)_3_ immunizations

(ccIZN36)_3_ was produced as previously described (7). Seven female Hartley guinea pigs from Charles River were immunized intramuscularly with (ccIZN36)_3_ at 0, 4, and 8 weeks. For each immunization, a total volume of 400 μL containing 100 μg (ccIZN36)_3_, 180 μg aluminum hydroxyphosphate sulfate, and 40 μg Iscomatrix (CSL Biotherapies) was evenly divided between two injection sites. Serum was collected 8 weeks before the first immunization and 3 weeks after each boost. Animal work was performed in accordance with IACUC 8119974780067.

## Acknowledgments

Research reported in this publication was supported by the National Institute of General Medical Sciences of the National Institutes of Health under award numbers T32GM007276 (to BNB) and 5T32GM007365 (to MVFI), NIH/NIAID contract #HHSN272201800004C (to CCL), the Bill and Melinda Gates Foundation (OPP1113682), the Virginia and D. K. Ludwig Fund for Cancer Research, and the Chan Zuckerberg Biohub (to PSK). The content is solely the responsibility of the authors and does not necessarily represent the official view of the National Institute of Health. BNB is supported by the NSF GRFP. We thank members of the Kim Lab for valuable discussions, Nielson Weng for consultation on statistics, and Drs. Lillian Cohn, Abigail Powell and Shaogeng Tang for their helpful feedback on this manuscript.

## References

1. D. C. Chan, S. Kim, HIV entry and its inhibition. Cell 93, 681–684 (1998).

2. D. M. Eckert, P. S. Kim, Mechanisms of viral membrane fusion and its inhibition. Annu. Rev. Biochem. 70, 777–810 (2001).

3. S. C. Harrison, Viral membrane fusion. Virology 479–480, 498–507 (2015).

4. J. M. Kilby, S. Hopkins, T. M. Venetta, B. Dimassimo, G. A. Cloud, J. Y. Lee, L. Alldredge, E. Hunter, et al., Potent suppression of HIV-1 replication in humans by T-20, a peptide inhibitor of gp41-mediated virus entry. Nat. Med. 4, 1302–1307 (1998).

5. J. LaBonte, J. Lebbos, P. Kirkpatrick, Enfuvirtide. Nat. Rev. Drug Discov. 2003 25 2, 345–346 (2003).

6. E. de Rosny, R. Vassell, P. T. Wingfield, C. T. Wild, C. D. Weiss, Peptides Corresponding to the Heptad Repeat Motifs in the Transmembrane Protein (gp41) of Human Immunodeficiency Virus Type 1 Elicit Antibodies to Receptor-Activated Conformations of the Envelope Glycoprotein. J. Virol. 75, 8859–8863 (2001).

7. E. Bianchi, J. G. Joyce, M. D. Miller, A. C. Finnefrock, X. Liang, M. Finotto, P. Ingallinella, P. McKenna, et al., Vaccination with peptide mimetics of the gp41 prehairpin fusion intermediate yields neutralizing antisera against HIV-1 isolates. Proc. Natl. Acad. Sci. U. S. A. 107, 10655–10660 (2010).

8. Z. Qi, C. Pan, H. Lu, Y. Shui, L. Li, X. Li, X. Xu, S. Liu, et al., A recombinant mimetics of the HIV-1 gp41 prehairpin fusion intermediate fused with human IgG Fc fragment elicits neutralizing antibody response in the vaccinated mice. Biochem. Biophys. Res. Commun. 398, 506–512 (2010).

9. J. D. Nelson, H. Kinkead, F. M. Brunel, D. Leaman, R. Jensen, J. M. Louis, T. Maruyama, C. A. Bewley, et al., Antibody elicited against the gp41 N-heptad repeat (NHR) coiled-coil can neutralize HIV-1 with modest potency but non-neutralizing antibodies also bind to NHR mimetics. Virology 377, 170–183 (2008).

10. J. M. Louis, I. Nesheiwat, L. C. Chang, G. M. Clore, C. A. Bewley, Covalent trimers of the internal N-terminal trimeric coiled-coil of gp41 and antibodies directed against them are potent inhibitors of HIV envelope-mediated cell fusion. J. Biol. Chem. 278, 20278–20285 (2003).

11. M. D. Miller, R. Geleziunas, E. Bianchi, S. Lennard, R. Hrin, H. Zhang, M. Lu, Z. An, et al., A human monoclonal antibody neutralizes diverse HIV-1 isolates by binding a critical gp41 epitope. Proc. Natl. Acad. Sci. U. S. A. 102, 14759–14764 (2005).

12. M. A. Luftig, M. Mattu, P. Di Giovine, R. Geleziunas, R. Hrin, G. Barbato, E. Bianchi, M. D. Miller, et al., Structural basis for HIV-1 neutralization by a gp41 fusion intermediate-directed antibody. Nat. Struct. Mol. Biol. 13, 740–747 (2006).

13. E. Gustchina, M. Li, J. M. Louis, D. Eric Anderson, J. Lloyd, C. Frisch, C. A. Bewley, A. Gustchina, et al., Structural basis of HIV-1 neutralization by affinity matured fabs directed against the internal trimeric coiled-coil of gp41. PLoS Pathog. 6 e1001182 (2010).

14. E. Gustchina, M. Li, R. Ghirlando, P. Schuck, J. M. Louis, J. Pierson, P. Rao, S. Subramaniam, et al., Complexes of neutralizing and non-neutralizing affinity matured fabs with a mimetic of the internal trimeric coiled-coil of HIV-1 gp41. PLoS One 8 e78187 (2013).

15. C. Sabin, D. Corti, V. Buzon, M. S. Seaman, D. L. Hulsik, A. Hinz, F. Vanzetta, G. Agatic, et al., Crystal structure and size-dependent neutralization properties of HK20, a human monoclonal antibody binding to the highly conserved heptad repeat 1 of gp41. PLoS Pathog. 6 e1001195 (2010).

16. M. S. Seaman, H. Janes, N. Hawkins, L. E. Grandpre, C. Devoy, A. Giri, R. T. Coffey, L. Harris, et al., Tiered categorization of a diverse panel of HIV-1 Env pseudoviruses for assessment of neutralizing antibodies. J. Virol. 84, 1439–52 (2010).

17. L. G. Perez, M. R. Costa, C. A. Todd, B. F. Haynes, D. C. Montefiori, Utilization of Immunoglobulin G Fc Receptors by Human Immunodeficiency Virus Type 1: a Specific Role for Antibodies against the Membrane-Proximal External Region of gp41. J. Virol. 83, 7397–7410 (2009).

18. L. G. Perez, S. Zolla-Pazner, D. C. Montefiori, Antibody-Dependent, Fc RI-Mediated Neutralization of HIV-1 in TZM-bl Cells Occurs Independently of Phagocytosis. J. Virol. 87, 5287–5290 (2013).

19. M. Montero, N. E. van Houten, X. Wang, J. K. Scott, The Membrane-Proximal External Region of the Human Immunodeficiency Virus Type 1 Envelope: Dominant Site of Antibody Neutralization and Target for Vaccine Design. Microbiol. Mol. Biol. Rev. 72, 54–84 (2008).

20. J. S. Gach, D. P. Leaman, M. B. Zwick, Targeting HIV-1 gp41 in Close Proximity to the Membrane Using Antibody and Other Molecules. Curr. Top. Med. Chem. 11, 2997–3021 (2011).

21. C. E. van der Poel, R. M. Spaapen, J. G. J. van de Winkel, J. H. W. Leusen, Functional Characteristics of the High Affinity IgG Receptor, FcγRI. J. Immunol. 186, 2699–2704 (2011).

22. S. Patterson, A. Rae, N. Hockey, J. Gilmour, F. Gotch, Plasmacytoid Dendritic Cells Are Highly Susceptible to Human Immunodeficiency Virus Type 1 Infection and Release Infectious Virus. J. Virol. 75, 6710–6713 (2001).

23. A. Smed-Sörensen, K. Loré, J. Vasudevan, M. K. Louder, J. Andersson, J. R. Mascola, A.-L. Spetz, R. A. Koup, Differential Susceptibility to Human Immunodeficiency Virus Type 1 Infection of Myeloid and Plasmacytoid Dendritic Cells. J. Virol. 79, 8861–8869 (2005).

24. R. Shen, H. E. Richter, R. H. Clements, L. Novak, K. Huff, D. Bimczok, S. Sankaran-Walters, S. Dandekar, et al., Macrophages in Vaginal but Not Intestinal Mucosa Are Monocyte-Like and Permissive to Human Immunodeficiency Virus Type 1 Infection. J. Virol. 83, 3258–3267 (2009).

25. Z. Kruize, N. A. Kootstra, The Role of Macrophages in HIV-1 Persistence and Pathogenesis. Front. Microbiol. 10 2828 (2019).

26. K. Loré, A. Smed-Sörensen, J. Vasudevan, J. R. Mascola, R. A. Koup, Myeloid and plasmacytoid dendritic cells transfer HIV-1 preferentially to antigen-specific CD4+ T cells. J. Exp. Med. 201, 2023–2033 (2005).

27. F. Groot, S. Welsch, Q. J. Sattentau, Efficient HIV-1 transmission from macrophages to T cells across transient virological synapses. Blood 111, 4660–4663 (2008).

28. K. Waki, E. O. Freed, Macrophages and cell-cell spread of HIV-1. Viruses 2, 1603–1620 (2010).

29. L. Bracq, M. Xie, S. Benichou, J. Bouchet, Mechanisms for cell-to-cell transmission of HIV-1. Front. Immunol. 9, 260 (2018).

30. A. I. Spira, P. A. Marx, B. K. Patterson, J. Mahoney, R. A. Koup, S. M. Wolinsky, D. D. Ho, Cellular targets of infection and route of viral dissemination after an intravaginal inoculation of simian immunodeficiency virus into rhesus macaques. J. Exp. Med. 183, 215–225 (1996).

31. J. Hu, M. B. Gardner, C. J. Miller, Simian Immunodeficiency Virus Rapidly Penetrates the Cervicovaginal Mucosa after Intravaginal Inoculation and Infects Intraepithelial Dendritic Cells. J. Virol. 74, 6087–6095 (2000).

32. M. Pope, A. T. Haase, Transmission, acute HIV-1 infection and the quest for strategies to prevent infection. Nat. Med. 9, 847–852 (2003).

33. S. G. Turville, J. J. Santos, I. Frank, P. U. Cameron, J. Wilkinson, M. Miranda-Saksena, J. Dable, H. Stõssel, et al., Immunodeficiency virus uptake, turnover, and 2-phase transfer in human dendritic cells. Blood 103, 2170–2179 (2004).

34. A. Martín-Moreno, M. A. Muñoz-Fernández, Dendritic cells, the double agent in the war against hiv-1. Front. Immunol. 10, 2485 (2019).

35. A. deCamp, P. Hraber, R. T. Bailer, M. S. Seaman, C. Ochsenbauer, J. Kappes, R. Gottardo, P. Edlefsen, et al., Global Panel of HIV-1 Env Reference Strains for Standardized Assessments of Vaccine-Elicited Neutralizing Antibodies. J. Virol. 88, 2489–2507 (2014).

36. E. Brocca-Cofano, K. McKinnon, T. Demberg, D. Venzon, R. Hidajat, P. Xiao, M. Daltabuit-Test, L. J. Patterson, et al., Vaccine-elicited SIV and HIV envelope-specific IgA and IgG memory B cells in rhesus macaque peripheral blood correlate with functional antibody responses and reduced viremia. Vaccine 29, 3310–3319 (2011).

37. A. Gonzalez-Quintela, R. Alende, F. Gude, J. Campos, J. Rey, L. M. Meijide, C. Fernandez-Merino, C. Vidal, Serum levels of immunoglobulins (IgG, IgA, IgM) in a general adult population and their relationship with alcohol consumption, smoking and common metabolic abnormalities. Clin. Exp. Immunol. 151, 42–50 (2008).

38. G. A. Roth, E. C. Gale, M. Alcántara-Hernández, W. Luo, E. Axpe, R. Verma, Q. Yin, A. C. Yu, et al., Injectable hydrogels for sustained co-delivery of subunit vaccines enhance humoral immunity. bioRxiv, 2020.05.26.117465 (2020).

39. Q. Gao, L. Bao, H. Mao, L. Wang, K. Xu, M. Yang, Y. Li, L. Zhu, et al., Development of an inactivated vaccine candidate for SARS-CoV-2. Science 369, 77–81 (2020).

40. K. Klein, R. S. Veazey, R. Warrier, P. Hraber, L. A. Doyle-Meyers, V. Buffa, H.-X. Liao, B. F. Haynes, et al., Neutralizing IgG at the Portal of Infection Mediates Protection against Vaginal Simian/Human Immunodeficiency Virus Challenge. J. Virol. 87, 11604–11616 (2013).

41. A. Pegu, B. Borate, Y. Huang, M. G. Pauthner, A. J. Hessell, B. Julg, N. A. Doria-Rose, S. D. Schmidt, et al., A Meta-analysis of Passive Immunization Studies Shows that Serum-Neutralizing Antibody Titer Associates with Protection against SHIV Challenge. Cell Host Microbe 26, 336–346.e3 (2019).

42. E. J. Platt, M. Bilska, S. L. Kozak, D. Kabat, D. C. Montefiori, Evidence that Ecotropic Murine Leukemia Virus Contamination in TZM-bl Cells Does Not Affect the Outcome of Neutralizing Antibody Assays with Human Immunodeficiency Virus Type 1. J. Virol. 83, 8289–8292 (2009).

43. Y. Takeuchi, M. O. McClure, M. Pizzato, Identification of Gammaretroviruses Constitutively Released from Cell Lines Used for Human Immunodeficiency Virus Research. J. Virol. 82, 12585–12588 (2008).

44. X. Wei, J. M. Decker, H. Liu, Z. Zhang, R. B. Arani, J. M. Kilby, M. S. Saag, X. Wu, et al., Emergence of resistant human immunodeficiency virus type 1 in patients receiving fusion inhibitor (T-20) monotherapy. Antimicrob. Agents Chemother. 46, 1896–1905 (2002).

45. C. A. Derdeyn, J. M. Decker, J. N. Sfakianos, X. Wu, W. A. O’Brien, L. Ratner, J. C. Kappes, G. M. Shaw, et al., Sensitivity of Human Immunodeficiency Virus Type 1 to the Fusion Inhibitor T-20 Is Modulated by Coreceptor Specificity Defined by the V3 Loop of gp120. J. Virol. 74, 8358–8367 (2000).

46. E. J. Platt, K. Wehrly, S. E. Kuhmann, B. Chesebro, D. Kabat, Effects of CCR5 and CD4 Cell Surface Concentrations on Infections by Macrophagetropic Isolates of Human Immunodeficiency Virus Type 1. J. Virol. 72, 2855–2864 (1998).

47. M. Li, J. F. Salazar-Gonzalez, C. A. Derdeyn, L. Morris, C. Williamson, J. E. Robinson, J. M. Decker, Y. Li, et al., Genetic and Neutralization Properties of Subtype C Human Immunodeficiency Virus Type 1 Molecular env Clones from Acute and Early Heterosexually Acquired Infections in Southern Africa. J. Virol. 80, 11776–11790 (2006).

48. P. Hraber, C. Rademeyer, C. Williamson, M. S. Seaman, R. Gottardo, H. Tang, K. Greene, H. Gao, et al., Panels of HIV-1 Subtype C Env Reference Strains for Standardized Neutralization Assessments. J. Virol. 91 e00991–17 (2017).

49. D. C. Montefiori, Measuring HIV neutralization in a luciferase reporter gene assay. Methods Mol. Biol. 485, 395–405 (2009).

50. M. Sarzotti-Kelsoe, R. T. Bailer, E. Turk, C. li Lin, M. Bilska, K. M. Greene, H. Gao, C. A. Todd, et al., Optimization and validation of the TZM-bl assay for standardized assessments of neutralizing antibodies against HIV-1. J. Immunol. Methods 409, 131–146 (2014).

51. D. A. Ozaki, H. Gao, C. A. Todd, K. M. Greene, D. C. Montefiori, M. Sarzotti-Kelsoe, International technology transfer of a GCLP-compliant HIV-1 neutralizing antibody assay for human clinical trials. PLoS One 7, e30963 (2012).

52. C. A. Todd, K. M. Greene, X. Yu, D. A. Ozaki, H. Gao, Y. Huang, M. Wang, G. Li, et al., Development and implementation of an international proficiency testing program for a neutralizing antibody assay for HIV-1 in TZM-bl cells. J. Immunol. Methods 375, 57–67 (2012).

